# Abandonment and restoration of Mediterranean meadows: long-term effects on a butterfly-plant network

**DOI:** 10.1101/2020.07.11.198499

**Authors:** Pau Colom, Anna Traveset, Constantí Stefanescu

## Abstract

Both the intensification and abandonment of traditional agricultural practices are known to be major threats to biodiversity worldwide, above all in industrialized countries. Although land abandonment in particular has a negative effect on the diversity of both plant and insect communities, few studies have ever analysed these two groups together and none has yet examined the effect on plant-insect interactions using a network approach. In view of the notable decline of pollinator insects reported in past decades, it is essential to understand how the structure of a plant-pollinator network changes during the ecological succession that occurs as traditionally managed habitats are abandoned, and to what extent this network is re-established when habitats are restored. We monitored a butterfly-plant network for 22 years in habitats where land abandonment and restoration have taken place and were able to compare restoration by grazing with restoration combining mowing and grazing. Abandonment leads to significant reductions in the cover of typical grasslands plants and, in turn, rapidly provokes changes in butterfly assemblages and plant interactions. Specifically, it caused a replacement of multivoltine by monovoltine species, increasing network specialization due to the great specificity in the interactions that monovoltine species established with plants. Changes in butterfly communities were also recorded in a nearby unaltered habitat due to the metapopulation structure of some species. A highly dynamic source-sink system was established between managed and unmanaged habitat patches, which ultimately allowed the metapopulations to persist. Restoration combining mowing and grazing promoted a quick return to the pre-abandonment situation in the butterfly community, and also increased generality and nestedness, two network descriptors that are known to enhance community stability.

## Introduction

Many of the most biodiverse systems of high ecological value in industrialized areas are the product of the development of traditional human agricultural practices (Di Giulio et al., 2001; Tscharntke et al., 2005; Kleijn et al., 2009, 2011; Blondel et al., 2010). However, many are now threatened by two contrasting phenomena, namely, agricultural intensification and abandonment (Donald et al., 2001; Briggs et al., 2005; Cramer et al., 2008; Kehoe et al., 2017; Zabel et al., 2019).

Numerous studies have found that grasslands managed by either mowing or grazing maintain a greater diversity of plant species than those that are abandoned or, by contrast, are subject to intense management (Poschlod & WallisDeVries, 2002; Pykälä et al., 2005). Disturbance caused by mowing or grazing causes an intermediate scenario that allows the least competitive species to survive, thereby favouring greater plant co-existence (Zobel et al., 1996). The literature describes an increase in insect species diversity at the beginning of the succession when meadows are abandoned or management intensity is reduced (Pöyry et al., 2006; Öckinger et al., 2006). The reason for this seems to be that taller but also structurally more diverse vegetation (i.e. increased diversity of turf heights) initially allows more diverse insect communities to thrive (Kruess & Tscharntke, 2002). Then, as ecological succession advances and shrub vegetation and trees encroach, diversity generally decreases, as has clearly been demonstrated in several butterfly studies (e.g. Balmer & Erhardt, 2000; Öckinger et al., 2006). In the Mediterranean region, where most species of butterflies show a strong preference for open habitats, afforestation resulting from the abandonment of traditional agricultural practices (Feranec et al., 2010) has been identified as one of the main factors driving population declines (Slancarova et al., 2016; Herrando et al., 2016; Ubach et al., 2019).

In conjunction, these studies have led to a broad consensus that, at least in Europe, a significant loss of biodiversity can be attributed to the abandonment of semi-natural meadows. This, in turn, has encouraged the recovery of such habitats via the restoration of traditional practices (Pykälä, 2003; Pöyry et al., 2005; Öckinger et al., 2006). However, restoring former management practices does not necessarily lead to an immediate return to the semi-natural state of the habitat prior to abandonment (Dover et al., 2011). Depending on the duration of the abandonment, it may take a long time for meadows to return to their former states (Rook et al., 2004). Moreover, the effects of the restoration on both plant and butterfly communities will depend on the type and the frequency of the management. Some studies suggest that the best results are obtained via grazing rather than mowing (Tälle et al., 2015, 2016), while others advocate an intermediate frequency (Bakker & Berendse, 1999; Watkinson & Ormerod, 2001). In addition, habitat recovery may depend on the type of grazer, since grazing by cows and horses in some cases has positive effects on plant and insect species richness, while grazing by sheep has been reported to have negative effects (Carvell, 2002; Öckinger et al., 2006).

In view of the notable decline in populations of insect pollinators in recent decades (Potts et al., 2010) and the valuable services they offer agriculture (Klein et al, 2007), important efforts are being made to understand how abandonment could affect this group. Specifically, the effects of abandonment and the restoration of grasslands have been studied in vascular plants (Meiners et al., 2001; Pruchniewicz, 2017; Uchida et al., 2018), insects (Erhardt, 1985; Marini et al., 2009; Dover et al., 2011) and in both groups simultaneously (Steffan-Dewenter & Leschke, 2003; Pöyry et al., 2006; Uchida & Ushimaru, 2015), although in the latter case, as far as we know, no network approach has been used. This approach is important because the presence of members of two interacting groups does not necessarily mean that their interactions are also restored. Given the importance of interactions between species in the functioning of ecosystems (Kremen, 2005), it is essential to understand not only the changes that occur in plant and insect communities independently of one another, but also the changes in the interactions occurring between these two groups. Also of importance are trends in network structures in the long term during periods of abandonment and restoration of semi-natural habitats (e.g. Olesen et al., 2008, 2011).

This work aimed to analyse the long-term effects of abandonment and the subsequent restoration of plant and butterfly communities in the same semi-natural meadows over a period of two decades. Specifically, we investigated how butterflies and their nectar plants respond to changes provoked by abandonment and, subsequently, to two different types of restoration. This study of both processes embraces analyses of habitat changes and their effects on species composition and population dynamics. Moreover, for the first time, we employed network analyses to understand the effects of abandonment and restoration on butterfly-plant interactions at community level.

## Material and methods

### Study site

Data were obtained from a system of traditionally managed meadows prone to flooding, where some of the effects of abandonment on both butterfly and plant communities have been reported in a previous study (Stefanescu et al., 2009). In the study area, a 1.1-km transect was established and divided into six sections (117–286-m long), each in a different meadow; sections were separated from each other by ditches or trees (see appendix A; Fig. 1).

Over a period of 22 years (1997–2018) these meadows underwent important changes in management practices. In the first two years, all six sections were managed traditionally, either by mowing (sections 1, 2, 5) or by a combination of grazing and mowing (sections 3, 4, 6). In 1999, sections 1–5 were abandoned (i.e. they were no longer grazed or mown), while section 6, which was managed as before (i.e. by mowing and grazing) throughout the whole study period, acted as a control (see appendix A: Fig. 2). Traditional management using grazing and mowing was restored in 2005 in all abandoned meadows (except section 1, which was only grazed). This type of management has continued up to the present day.

### Data sampling

The butterfly populations of this area have been monitored since 1997 within the framework of the Catalan Butterfly Monitoring Scheme (<www.catalanbms.org>). Butterfly individuals were counted following a standardized methodology: from the first week of March to the last week of September weekly samplings along a walked transect were taken at distances of 2.5 m on both the sides and 5 m ahead of the recorder (Pollard & Yates, 1993). Based on these counts, annual indices of relative abundances per species were calculated to evaluate population trends during the study period. Abundance values were standardized per 100 m of section length. The interactions and their frequency (i.e. number of floral visits) between butterflies and their nectar plants were also recorded. These records only include butterflies that were actually seen to feed on nectar with their proboscis clearly extended. All interaction data were arranged in annual bipartite matrices.

In 2000 for the first time, plant communities were characterized following the CORINE Land Cover manual and then repeated every six years. This vegetation monitoring thus provides information regarding which plant communities were dominant in the meadows in the first year after the abandonment (2000), in the year after management recovery (2006), and in two subsequent monitoring periods (2012 and 2018).

### Data analysis

#### Ecological descriptors

The combination of data on butterflies and flowering plants was used to characterize ecological changes during the study period. The following five ecological descriptors (hereafter EDs) were calculated annually for each section: (1) Butterfly abundance; (2) Butterfly species richness; (3) Butterfly diversity (calculated using the Shannon-Wiener index); (4) Flower visits (i.e. total number of flower visits); and (5) Plant species richness (i.e. number of species of flowering plants visited by butterflies). Trends over time in the EDs in the different sections and periods were analysed using linear models. Differences in the ED trends between managed (control section #6) and abandoned (#1–5) sections during the abandonment period, and between the two types of restoration (only-grazed section 1 vs. mown/grazed sections 2-5) were analysed using Generalised Linear Models (GLM).

#### Butterfly and nectar-plant species composition

For the analysis of plant species composition (i.e. flowering plants visited by butterflies), all sections were pooled due to the limited number of recorded flower-butterfly interactions in some years at section level. Bray-Curtis dissimilarity indices were used to measure changes in composition between abandonment and management periods. The same indices were used to analyse changes in the composition of butterfly communities by comparing each section and year with the initial community (i.e. in 1997, the first year). Temporal trends in dissimilarity values were then calculated using linear models for the three different periods of the study, including as a reference value the year previous to the change of management type: (1) abandonment of sections 1–5 (1998–2004); (2) recovery of traditional management in sections 1–5 (2004–2018); and (3) the whole study period (1997-2018). PERMANOVA analysis and NMDS plots were also conducted to test for possible differences in species composition between periods and sections. SIMPER analysis (Clarke, 1993) was additionally used to identify the species that contributed most to the total dissimilarity between abandonment and restoration periods.

#### Butterfly population trends related to ecological and life history traits

We tested whether butterfly population trends (measured as changes in the annual indices of relative abundance) could be explained by species traits during the two study periods (abandonment and management). The selected ecological and life-history traits of the butterflies were: (a) wing length (wing span, from García-Barros et al., 2013); (b) mobility according to a categorical index with five classes (0 = populations showing very little dispersal; 1 = closed populations and more frequent dispersal; 2 = closed populations and very frequent dispersal; 3 = open populations and non-directional dispersal; and 4 = open populations and directional migration; see Stefanescu et al., 2005, 2009); (c) overwintering stage (egg, larva, pupa, adult, or no overwintering stage); (d) host-plant specialization as a larva (i.e. monophagous, oligophagous or polyphagous); (e) voltinism (i.e. number of generations per year: monovoltine, bivoltine or multivoltine). Life-history data were extracted from García-Barros et al. (2013), Vila et al. (2018) and personal observations by one of the authors (CS).

#### Network analysis

To evaluate the temporal dynamics of the butterfly-plant interactions over the study period, we calculated different network-level indices commonly used in network analysis with the *bipartite* package in R version 3.6.3 (Dormann et al., 2009). Specifically, specialization index (*H*_*2*_’) modularity and the nestedness (WNOF) of the network were obtained annually for the whole set of abandoned meadows (i.e. excluding the control section); sections were pooled due to the low number of visits per section per year. Generalized Additive Models (GAM) were used to analyse the trends of indices during the abandonment-restoration succession.

## Results

### Habitat changes related to meadow management

Despite the lack of data on plant community composition prior to abandonment, important changes were observed between 2000 and 2006 as a result of the cessation of grazing and mowing. By 2000 (i.e. one year after abandonment), Mediterranean grasslands with *Gaudinia fragilis* and *Brachypodium phoenicoides* dominated the entire transect. Other species such as *Euphorbia serrata* and *Galium lucidum*, and typical wetland species such as *Scirpus holoschoenus*, were also abundant. In the period 2000–2006, however, Mediterranean grassland cover fell by 36±18% in the abandoned sections. Once the traditional management was restored, this habitat type increased again (24±26%) in those sections where mowing and grazing were combined. By contrast, the grassland community continued to decline (12% fall in 2006–2012) in the only-grazed section until it had completely disappeared by 2018 (see appendix A: Fig. 3). In this section, the grassland community was almost completely replaced by riparian woodland (mainly *Fraxinus angustifolia* and *Ulmus minor*; 50% of the coverage in 2018). Such notable increase in the riparian woodland coverage only occurred in one meadow restored by both mowing and grazing (#4), where a stand of *Populus alba* established itself (40% of coverage after abandonment and 60% after restoration). Although management never ceased in the control section (#6), this meadow became severely ruderalized (11% in 2000 vs. 90% in 2018 of ruderal habitat coverage), probably due to overgrazing by horses (see appendix A; Fig. 3). Indeed, the annual number of episodes of grazing and mowing in the control section was greater than in any other section (3.36 vs. 2.2 ± 0.3) (see appendix A: Fig. 2).

### Trends in ecological descriptors

During the whole of the study period the only-grazed section (# 1) was the only meadow that showed significant temporal trends in all EDs (Table 1; see appendix A: Fig. 4). Butterfly diversity declined significantly during the abandonment period in this section, and butterfly abundance, butterfly species richness, total number of flower visits and plant species richness showed negative but non-significant trends. After management was restored in 2005, differences between the two types of restoration became patent since the richness and diversity of butterflies and the number of flower visits decreased in the only-grazed section (#1) but increased or remained stable in the mown-grazed sections (#2-5). Differences between managed and abandoned sections were observed only in the number of visited plants, for which only the control showed a great increase in the richness of visited plants during abandonment. Despite being constantly managed over all the years of the study, the control section showed significant negative trends in butterfly richness and diversity for the whole period. During the abandonment of the other sections butterfly and plant abundance and richness of visited plants increased in the control section, although the same EDs decreased significantly once management was restored.

**Table 1.** Temporal trends in all the ecological descriptors in the analysed periods. Beta coefficients and *P* values are given. *P* values for the Generalised Linear Model comparisons between the trends in the different treatments (abandoned vs. managed and grazing vs. grazing + mowing) are shown in the GLM * rows.

|  | Study period<br>(1997–2018) |  | Abandonment<br>(1998–2004) |  | Restoration<br>(2004–2018) |  |
| --- | --- | --- | --- | --- | --- | --- |
|  |  |  | Abandoned | Managed | Grazing | Graz. + mown |
| **Butterfly abundance**<br>(Number of individuals/100m) | S1 | $\beta = -7.759$<br>$P < 0.001$ | $\beta = -5.85$<br>$P = 0.528$ | | $\beta = -4.013$<br>$P = 0.229$ | |
| | S2 | $\beta = -0.322$<br>$P = 0.906$ | $\beta = 29.73$<br>$P = 0.087$ | | | $\beta = -3.944$<br>$P = 0.382$ |
| | S3 | $\beta = -0.689$<br>$P = 0.753$ | $\beta = 29.42$<br>$P = 0.051$ | | | $\beta = -2.424$<br>$P = 0.459$ |
| | S4 | $\beta = -4.686$<br>$P = 0.037$ | $\beta = 21.63$<br>$P = 0.125$ | | | $\beta = -0.209$<br>$P = 0.923$ |
| | S5 | $\beta = 1.272$<br>$P = 0.643$ | $\beta = -6.67$<br>$P = 0.415$ | | | $\beta = 3.434$<br>$P = 0.552$ |
| | S6 | $\beta = -3.272$<br>$P = 0.302$ | | $\beta = 33.65$<br>$P = 0.004$ | | $\beta = -13.88$<br>$P = 0.019$ |
| | GLM* | | $F = 3.982; P = 0.074$ | | $F = 1.18; P = 0.287$ | |
| **Butterfly richness**<br>(Number of butterfly species) | S1 | $\beta = -0.579$<br>$P < 0.001$ | $\beta = -0.571$<br>$P = 0.263$ | | $-0.489$<br>$P = 0.022$ | |
| | S2 | $\beta = 0.183$<br>$P = 0.092$ | $\beta = 0.285$<br>$P = 0.715$ | | | $\beta = 0.111$<br>$P = 0.539$ |
| | S3 | $\beta = 0.046$<br>$P = 0.711$ | $\beta = 1.643$<br>$P = 0.074$ | | | $\beta = -0.075$<br>$P = 0.72$ |
| | S4 | $\beta = -0.327$<br>$P = 0.008$ | $\beta = -0.286$<br>$P = 0.740$ | | | $\beta = 0.096$<br>$P = 0.518$ |
| | S5 | $\beta = -0.208$<br>$P = 0.047$ | $\beta = 0.285$<br>$P = 0.363$ | | | $\beta = 0.079$<br>$P = 0.669$ |
| | S6 | $\beta = -0.254$<br>$P = 0.031$ | | $\beta = 0.893$<br>$P = 0.153$ | | $\beta = -0.371$<br>$P = 0.055$ |
| | GLM* | | $F = 0.844; P = 0.38$ | | $F = 6.446; P = 0.017$ | |
| **Butterfly diversity**<br>(Shannon diversity index of butterflies) | S1 | $\beta = -0.043$<br>$P < 0.001$ | $\beta = -0.077$<br>$P = 0.004$ | | $\beta = -0.048$<br>$P = 0.003$ | |
| | S2 | $\beta = 0.018$<br>$P = 0.005$ | $\beta = -0.047$<br>$P = 0.132$ | | | $\beta = 0.031$<br>$P = 0.021$ |
| | S3 | $\beta = 0.019$<br>$P = 0.008$ | $\beta = 0.069$<br>$P = 0.126$ | | | $\beta = 0.012$<br>$P = 0.303$ |
| | S4 | $\beta = -0.021$ | $\beta = -0.047$ | | | $\beta = 0.013$ |
| | | **$P = 0.032$** | $P = 0.294$ | | | $P = 0.391$ |
| | S5 | **$\beta = -0.034$**<br>**$P = 0.028$** | $\beta = 0.047$<br>$P = 0.465$ | | | $\beta = -0.019$<br>$P = 0.459$ |
| | S6 | **$\beta = -0.031$**<br>**$P = 0.002$** | | $\beta = -0.009$<br>$P = 0.616$ | | $\beta = -0.022$<br>$P = 0.244$ |
| | GLM* | | $F = 0.002; P = 0.963$ | | **$F = 12.54; P = 0.003$** | |
| **Flower visits**<br>(Total number of plants visited by butterflies) | S1 | **-7.47**<br>**$P < 0.001$** | $\beta = -3.25$<br>$P = 0.344$ | | **$\beta = -4.854$**<br>**$P = 0.014$** | |
| | S2 | $\beta = -0.016$<br>$P = 0.992$ | $\beta = 20.21$<br>$P = 0.109$ | | | $\beta = 0.221$<br>$P = 0.924$ |
| | S3 | $\beta = 0.638$<br>$P = 0.416$ | $\beta = 8.786$<br>$P = 0.123$ | | | $\beta = 1.989$<br>$P = 0.076$ |
| | S4 | $\beta = 0.189$<br>$P = 0.424$ | **$\beta = 1.857$**<br>**$P = 0.035$** | | | **$\beta = 0.901$**<br>**$P = 0.044$** |
| | S5 | $\beta = 0.998$<br>$P = 0.433$ | $\beta = 4.714$<br>$P = 0.356$ | | | $\beta = 1.296$<br>$P = 0.62$ |
| | S6 | $\beta = -0.937$<br>$P = 0.278$ | | **$\beta = 10.39$**<br>**$P = 0.006$** | | **$\beta = -3.914$**<br>**$P = 0.015$** |
| | GLM* | | $F = 1.094; P = 0.32$ | | **$F = 7.727; P = 0.01$** | |
| **Plant richness**<br>(Number of plant species visited by butterflies) | S1 | **$\beta = -0.028$**<br>**$P = 0.011$** | $\beta = -0.714$<br>$P = 0.13$ | | $\beta = -0.278$<br>$P = 0.16$ | |
| | S2 | **$\beta = 0.199$**<br>**$P = 0.017$** | $\beta = 0.286$<br>$P = 0.535$ | | | $\beta = 0.203$<br>$P = 0.194$ |
| | S3 | $\beta = 0.117$<br>$P = 0.274$ | $\beta = 0.643$<br>$P = 0.153$ | | | $\beta = -0.157$<br>$P = 0.449$ |
| | S4 | $\beta = -0.053$<br>$P = 0.424$ | $\beta = 0.25$<br>$P = 0.392$ | | | $\beta = 0.05$<br>$P = 0.701$ |
| | S5 | $\beta = 0.111$<br>$P = 0.07$ | $\beta = -0.25$<br>$P = 0.548$ | | | **$\beta = 0.304$**<br>**$P = 0.001$** |
| | S6 | $\beta = -0.076$<br>$P = 0.401$ | | **$\beta = 1.143$**<br>**$P = 0.032$** | | **$\beta = -0.425$**<br>**$P = 0.008$** |
| | GLM* | | **$F = 5.109; P = 0.047$** | | $F = 3.203; P = 0.085$ | |

### Changes in butterfly and flowering plant community structures

During the 22 study years significant changes occurred in butterfly communities with respect to the control year (1997). Such changes were observed not only in the only-grazed section (#1) but also in the control section (#6), although the trend was much more marked in the former (*R*^*2*^ = 0.73 vs. *R*^*2*^ = 0.18, respectively) (Fig. 1). During the abandonment period, butterfly communities changed significantly in all sections (Fig. 1, appendix A: Table 1). However, after management was restored, the only-grazed section (#1) was the only one in which butterfly communities became increasingly more dissimilar relative to the control year. By contrast, in the mowing-grazing sections the dissimilitude values decreased after management was restored, indicating a return to the initial state of the communities prior to abandonment. The temporal trends in butterfly and flowering plant composition were confirmed by PERMANOVA and NMDS analyses (Fig. 2, appendix A: Table 2). Butterfly communities during the abandonment period differed significantly from those during the restoration period (Fig. 2). These analyses further showed differences between the two types of restoration, as only the grazing-restored section (#1) showed significant differences between the prior to abandonment, abandonment and restoration periods. By contrast, the mowing-grazing sections (#2-5) did not show any significant differences between the restoration period and prior to abandonment, indicating that this type of combined management was an effective restoration technique.

**Figure 1.**
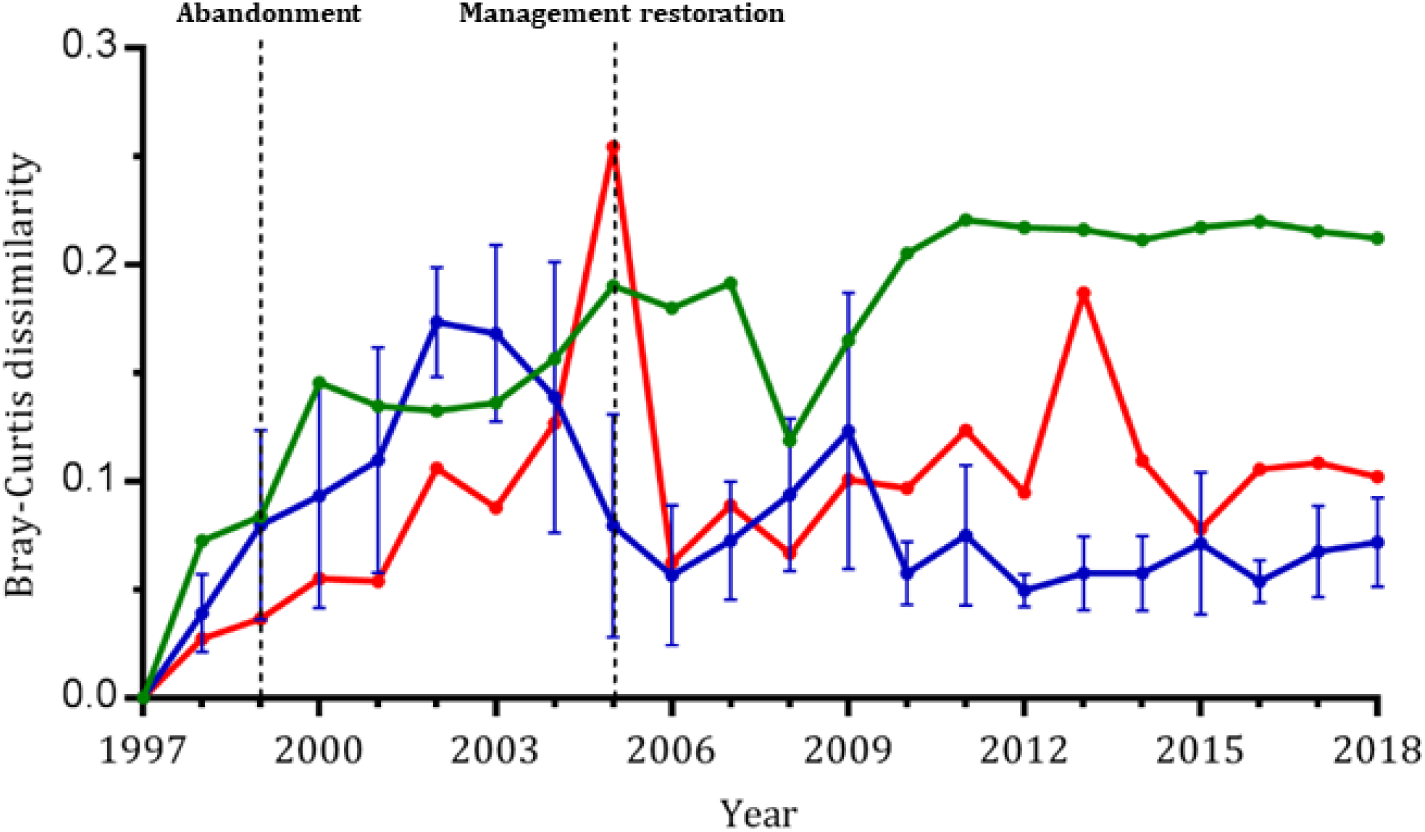
Trends in dissimilarity values with respect to the first year of monitoring (1997) for the butterfly communities. In green: section 1; in blue: sections 2–5 (with standard deviation represented by bars); in red: section 6.

**Figure 2.**
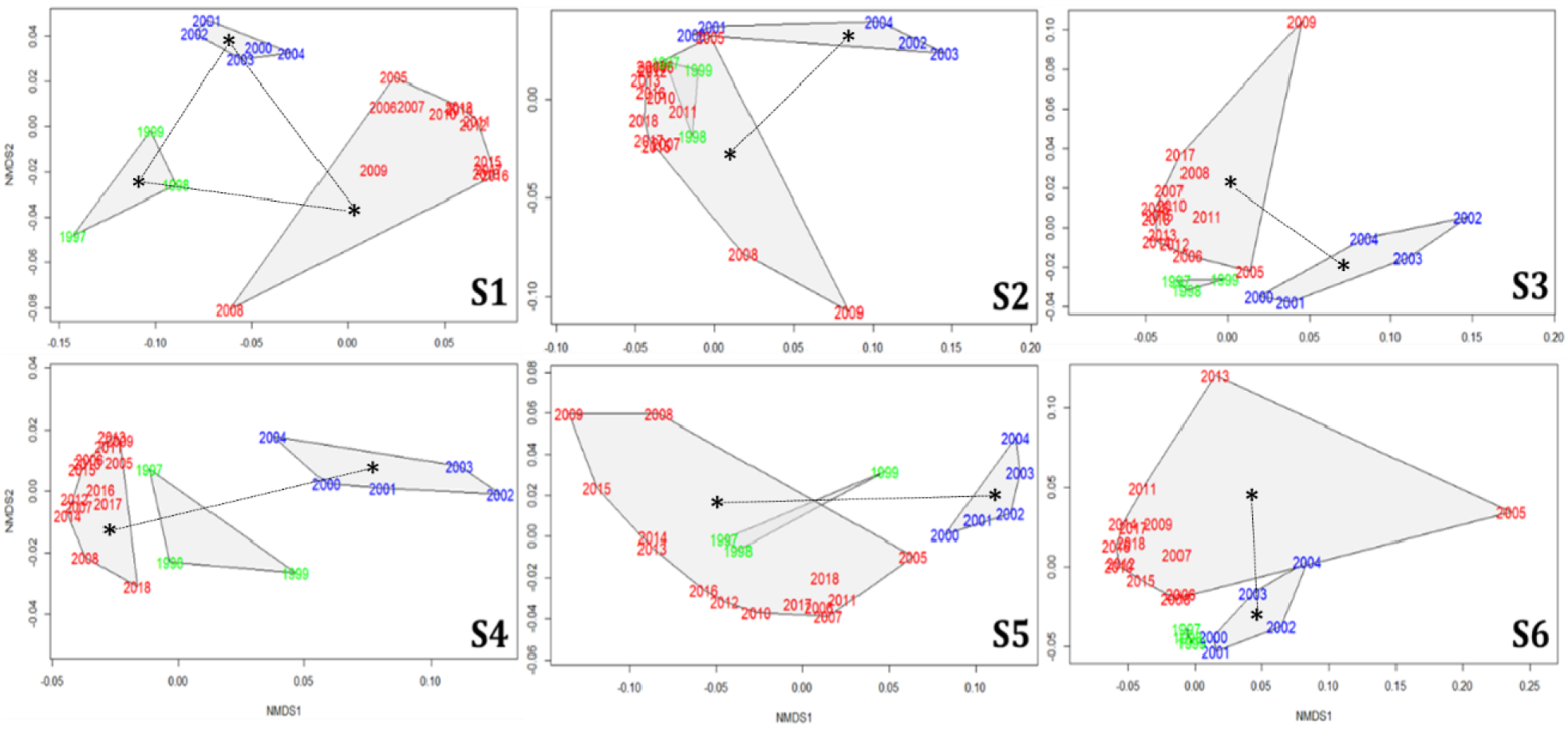
Non-Metric Multidimensional Scaling (NMDS) plots for each section of the transect. The different periods of analysis are represented in colours: green: before abandonment; blue: abandonment period; red: after management was restored. Dotted lines and asterisks represent significant differences (*P* < 0.05 in PERMANOVA analysis) between periods. * 1999 was included in the period before abandonment as butterfly communities were still very similar to the initial situation due to the inertia in changes in plant composition in the first year after abandonment.

The analyses also revealed significant differences in flowering plant composition between the abandonment and management periods at transect level (*F* = 11.5; *P* = 0.009). The species most visited by butterflies in the abandonment period were *Cirsium* spp., *Rubus* spp. and *Mentha suavolens*, which dominated the whole of the butterfly-plant interactions recorded during that period (see appendix A; Fig. 5). Once management was restored, however, the number of visits to these plants fell dramatically (except for *Rubus* spp., which maintained a large number of visits throughout the whole study period), while the number of visits to *Lotus corniculatus* and several species of *Trifolium* increased, despite the severe fall in number of visits during the abandonment period.

### Butterfly ecological traits related to management

Voltinism was the only ecological trait that predicted butterfly population trends in the abandoned sections. This trait explained the observed population trends during both the abandonment and the management periods, albeit with opposing effects (Fig. 3). All monovoltine species experienced positive trends when the meadows were abandoned but negative trends once management was restored. Bivoltine and multivoltine species, on the other hand, showed fairly variable trends in both periods, although multivoltine species tended to benefit from grazing and mowing as indicated by mostly positive trends during the restoration period (Fig. 3).

**Figure 3.**
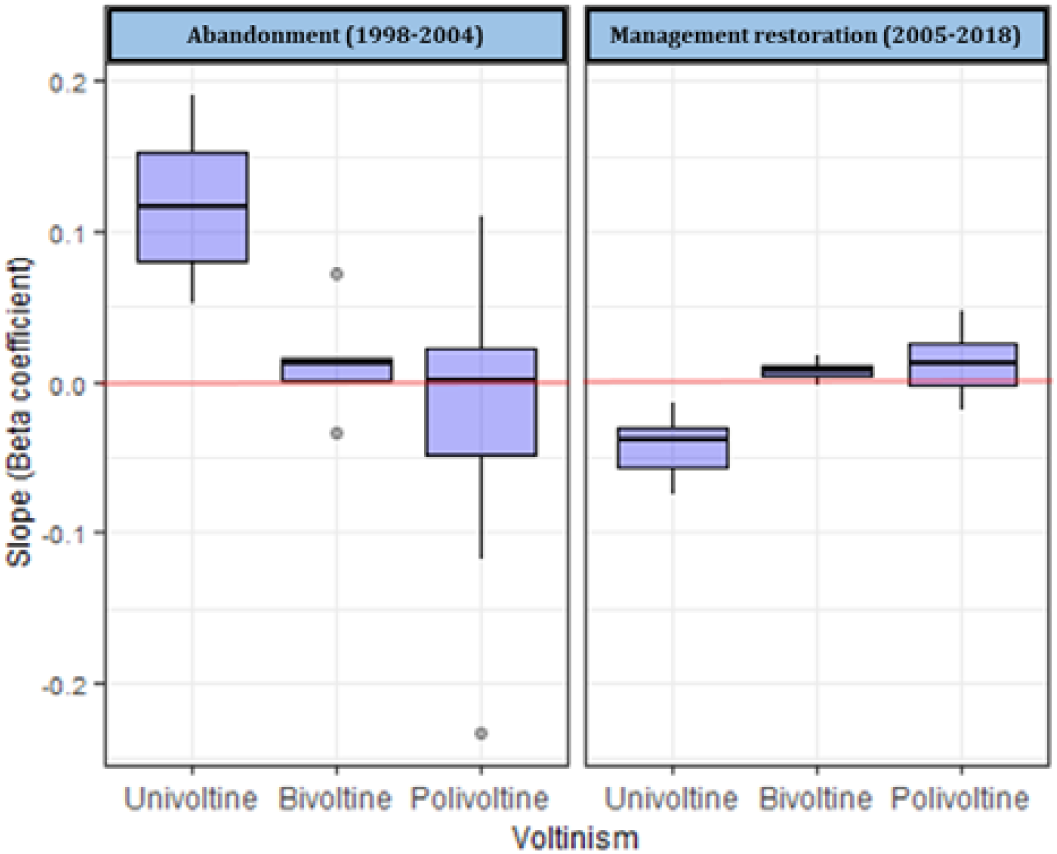
Population trends in the butterfly species in the abandoned sections (1–5) in relation to voltinism (number of generations per year). Population trends in the different species are represented as the slope values of the linear regression models.

### Butterfly-plant interaction trends

A total of 45 species of butterflies and 65 species of nectar plants were recorded and 17% of all possible interactions between these groups were detected. The species present in each annual network were highly variable and showed low persistence (i.e. number of years present) in many cases. Persistence was lower in plants (7.6 ± 6.7 years) than in butterflies (13 ± 8.8 years) (Wilcoxon rank test: W = 2439.5; *P* < 0.001). The high turnover of species indicates great variability in the interactions occurring between butterflies and plants from year to year, which translates into a variable network structure over time. Despite the great annual fluctuation in network parameters, results were consistent with a temporal trend of linear decrease in network specialization (*H*_*2*_’) (R^2^ adjusted = 0.34) (Fig. 4). However, when the only-grazed section (# 1) was excluded, this trend fitted a polynomial function better (adjusted R^2^ = 0.57), which showed how network specialization increased during the abandonment period and diminished again after management was restored. Modularity significantly decreased over time across all sections, although this relationship was not significant if the only-grazed section was excluded. The mowing-grazing restoration showed an overall increase in nestedness, although its trend was only marginally significant.

**Figure 4.**
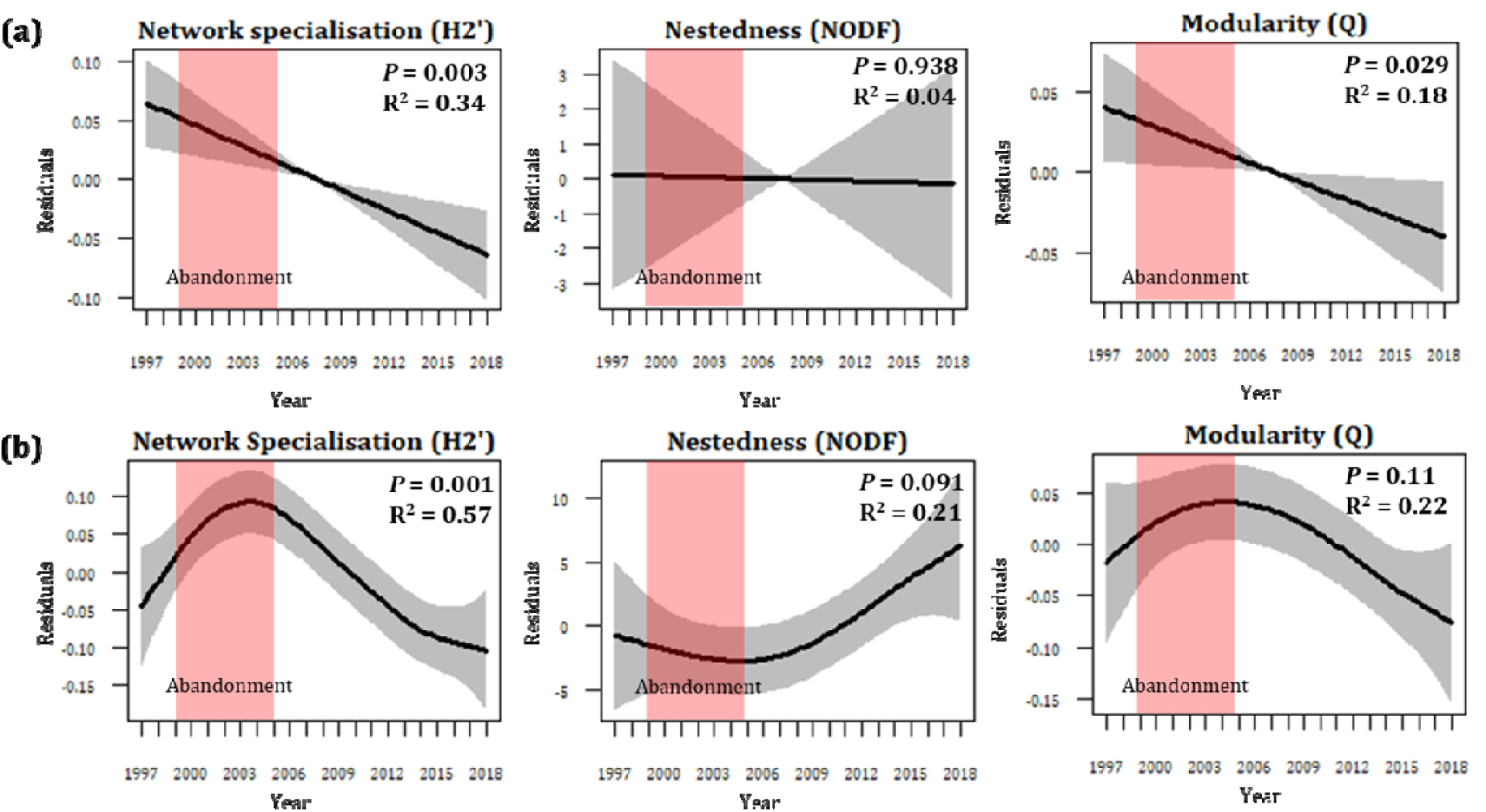
Network structure dynamics in the abandoned meadows. Network indices trends for (a) all abandoned meadows – i.e. sections 1–5 of the transect - and (b) abandoned sections that were restored by mowing and grazing – i.e. sections 2–5 of the transect. P-values and adjusted R-squared of the GAM models are shown.

## Discussion

A vast amount of literature shows how rapidly butterfly populations respond to habitat changes of different kinds (e.g. Thomas, 1991; Dennis, 2010). Evidence for such responses has rapidly accumulated over past decades in many European countries thanks to the establishment of butterfly monitoring schemes and the recognition of butterflies as a good bioindicator group (Thomas, 2005). The present work illustrates not only how butterfly populations respond to changes in habitats but also that notable changes occur in the interactions established between plants and adult butterflies.

### Impact of abandonment and restoration on butterflies and plant interactions

The decline in some butterfly species and their interactions with many plants was probably linked to the loss of nearly half of the grassland coverage after the meadows were abandoned. Restoration enabled these plant communities to recover significantly when mowing and grazing were combined (see appendix A: Fig. 3). Otherwise, pasturing alone proved to be insufficient in one of the meadows (#1), where grassland communities continued to decline until they completely disappeared by 2018. Therefore, mowing may be necessary for the conservation of typical Mediterranean meadows, which in this protected area have been identified as the most valuable habitat for butterflies (Stefanescu et al., 2005).

Butterfly community analysis over time confirms that changes in habitat lead to rapid modifications in butterfly assemblages. Thus, once the meadows were abandoned, butterfly communities underwent dramatic rearrangements, with a few monovoltine grass-feeder species experiencing population explosions (e.g. *Melanagia lachesis, Pyronia cecilia* and *Pyronia tithonus*, see appendix A: Fig. 6). By contrast, the populations of some multivoltine legume-feeders collapsed (e.g. *Plebejus argus* and *Polyommatus icarus*). Interestingly, changes in the populations of these species were also noticed in the control section (#6) that was managed throughout the whole study period, although these trends were in the opposite direction. As the meadows were abandoned, both butterfly abundances and visits to flowering plants substantially increased in the control section. Because the habitat remained essentially the same in the control section during this period, population increases of multivoltine species were probably related to the forced dispersal of populations from the nearby abandoned meadows. This is exemplified by *Plebejus argus*, whose numbers increased dramatically in the control section (up to an extraordinary density of five individuals/m in 2005) coinciding with the collapse of its populations in the abandoned sections. Therefore, population fluctuations are not only related to changes in the habitat where they are recorded but are also likely to be affected by a wider range of habitats where they are connected to other subpopulations (Keymer et al., 2000; Johst et al., 2002) in a metapopulation structure (Thomas & Harrison, 1992; Hanski & Thomas, 1994; Leweis et al., 1997). The temporal trends show how a meadow representing a sink habitat for *P. argus* at the beginning of the study became the only stronghold for its vanishing populations among the abandoned and deteriorating meadows. Moreover, once the habitat in the meadows improved following the restoration of management, the single meadow harbouring a population (i.e. the former sink) of this butterfly became a source (Pulliam, 1988) from which new habitats were re-colonized. This pattern highlights the importance of conserving networks of well-connected patches for habitat specialists, as has been highlighted by many theoretical and empirical studies (e.g. Hanski, 1999). This highly dynamic system allowed for a rapid recovery of collapsing populations when mowing and grazing restarted in the abandoned meadows harbouring the original populations.

The complexity of the management techniques required to reach ideal conditions for plant and butterfly communities is a recurrent theme in butterfly conservation (Settele et al., 2009). In this context, the control meadow where management continued throughout the study period showed some of the worst trends. This is likely due to the great disturbance (i.e. large number of grazing and mowing episodes), which ultimately affected butterfly populations and interactions with plants. The ruderalization of plant communities was linked to a reduction in the number and diversity of flower resources, which led to lower abundances in butterfly and probably other pollinator communities (Scheper et al., 2014). The absence of negative trends in butterfly richness and diversity suggests that the loss of available nectar resources led to a reduction in butterfly abundance and not *vice versa*.

### Butterfly life-history traits predict species’ responses to environmental succession

Certain previous studies have attempted to explain trends in butterfly populations in abandoned grasslands by examining species traits (Steffan-Dewenter & Tscharntke, 1997; Sanford, 2002; Stefanescu et al., 2009). Kithara et al. (2000) reported that species richness declined more in specialists than in generalists along a gradient of increasing disturbance when specialization was measured in terms of voltinism and host-plant specialization. Likewise, Pöyry et al. (2006) observed that the abandonment of grasslands benefitted generalist herbivores, while low-intensity management was more beneficial to specialists. In our study system, Stefanescu et al. (2009) failed to observe differences in host-plant specialization but did detect an increase in seasonal specialization (i.e. decrease in voltinism) of the communities in accordance to the r/k species concept (Pianka, 1970). In a habitat with recurrent disturbance (i.e. mowing and/or grazing) the species that will dominate the community will be those with high reproductive rates (Brown & Southwood, 1983; Brown, 1985). By contrast, species with longer developmental times and, therefore, with fewer annual generations will benefit from the abandonment of management practices (i.e. absence of disturbance). Our results, added to the previous study by Stefanescu et al. (2009) using data from another 14 years of management restoration, confirm that voltinism is indeed the best life-history trait for predicting population trends affected by managing practices in Mediterranean meadows.

### How does network structure change as a result of meadow management?

Our work also revealed interesting changes in the butterfly-plant network structure as a consequence of habitat management. The non-linearity of long-term trends in network parameters makes sense if we consider that management practices changed twice during the study period (Fig. 4). The increase in network specialization (*H*_*2*_’) when there was a lack of management suggests greater feeding selectivity by butterflies in this period. Butterflies are commonly regarded as generalist nectar-feeders, the level of specialization of species being more related to the length of the flight period than to their evolutionary history (Stefanescu & Traveset. 2009; Olesen et al., 2011). Monovoltine species will only be able to interact with those plants that are in bloom during their short flight period, whereas species with multiple generations can potentially interact with a greater number of plants. Therefore, the turnover of species that occurred during meadow abandonment (i.e. the substitution of multivoltine by monovoltine species) could explain the increasing specialization of the network. In other words, the greater specialization of the network was probably not due to a change in species’ behaviour but to population changes and species turnover. Moreover, although neither the total number of visits nor the diversity of the plants visited significantly changed in this period (Table 1), the number of butterfly visits some opportunistic plants (e.g. *Cirsium* spp., *Rubus* spp. and *Mentha suavolens*) received increased markedly (see appendix A: Fig. 5). The dominance of these species could have reduced the likelihood that butterflies would interact with other species. Previous studies have reported a relative constancy in the nestedness pattern in plant-pollinator networks subject to notable annual fluctuations in the identity of the species in the network (Alarcón et al., 2008; Petanidou et al., 2008). Nevertheless, we found a slight tendency for nestedness to increase in the meadows that were restored (*P* = 0.091, R^2^ = 0.21). This pattern could amplified given the increasing complexity of the network (i.e. number of interactions) as more abundant butterfly populations will predictably visit more nectar plant species (Bascompte et al., 2003). Both an increase in generality (i.e. reducing network specialization) and network nestedness could enhance the stability and functional redundancy of communities (Okuyama & Holland, 2008; Kaiser-Bunbury et al., 2017). This could be especially important in a context of global change, where episodes of extreme aridity are likely to threaten Mediterranean butterfly populations (Herrando et al., 2019). Therefore, the restoration of traditional management in meadows could increase ecosystem resilience in the face of future climate disturbance (Walker, 1995). In this context, we consider that restoration efforts in semi-natural habitats should be focused not only on species but also on their interactions (Tylianakis et al., 2010; Valiente Banuet et al., 2015; Kaiser-Bunbury and Blütghen, 2015).

## Conclusions

The abandonment of Mediterranean meadows led to significant reductions in the cover of typical grassland plants and, in turn, caused rapid changes to occur in butterfly assemblages. Such changes were recorded not only in meadows undergoing vegetation encroachment but also in nearby unaltered habitats due to the metapopulation structure of some butterfly species. A highly dynamic source-sink system was then established between managed and unmanaged habitat patches, ultimately allowing for metapopulation persistence. In addition, our data show that management restoration combining mowing and grazing can promote a quick return in the butterfly community to the pre-abandonment situation. However, insufficient management pressure (only-grazed section 1) or, conversely, excessive grazing and mowing pressure (control section 6) did not permit a proper recovery and led, instead, to a progressive decline in diversity. Interesting temporal trends in the butterfly-plant network structure paralleling habitat changes were also detected. Both interaction generalisation and nestedness decreased when meadows were abandoned but increased again once habitat was restored by combining mowing and grazing. These results suggest that effective meadow management not only helps maintain a richer butterfly community but also increases functional redundancy and network stability. Our work highlights the importance of maintaining traditional management practices in these semi-natural meadows as an effective way of preserving their highly diverse communities and the stability of the whole butterfly-plant network.

## Acknowledgements

We wish to thank the staff of the Aiguamolls de l’Empordà Natural Park for their support during these years. Thanks too are due to Ferran Páramo and Andreu Ubach for their technical support, Amparo Lázaro for her assistance in some of the statistical analyses and Pedro Bergamo for his useful comments and suggestions in an advanced version of the manuscript. The Butterfly Monitoring Scheme in Catalonia is funded by the Departament de Territori i Sostenibilitat de la Generalitat de Catalunya. PC was funded by a PhD fellowship from the Govern de les Illes Balears (FPI-CAIB-2018). This work is part of project GLC2017-88122-P financed by the Spanish Ministry of Science and Innovation to AT.

## Appendix A. Supplementary data

**Table A1.**
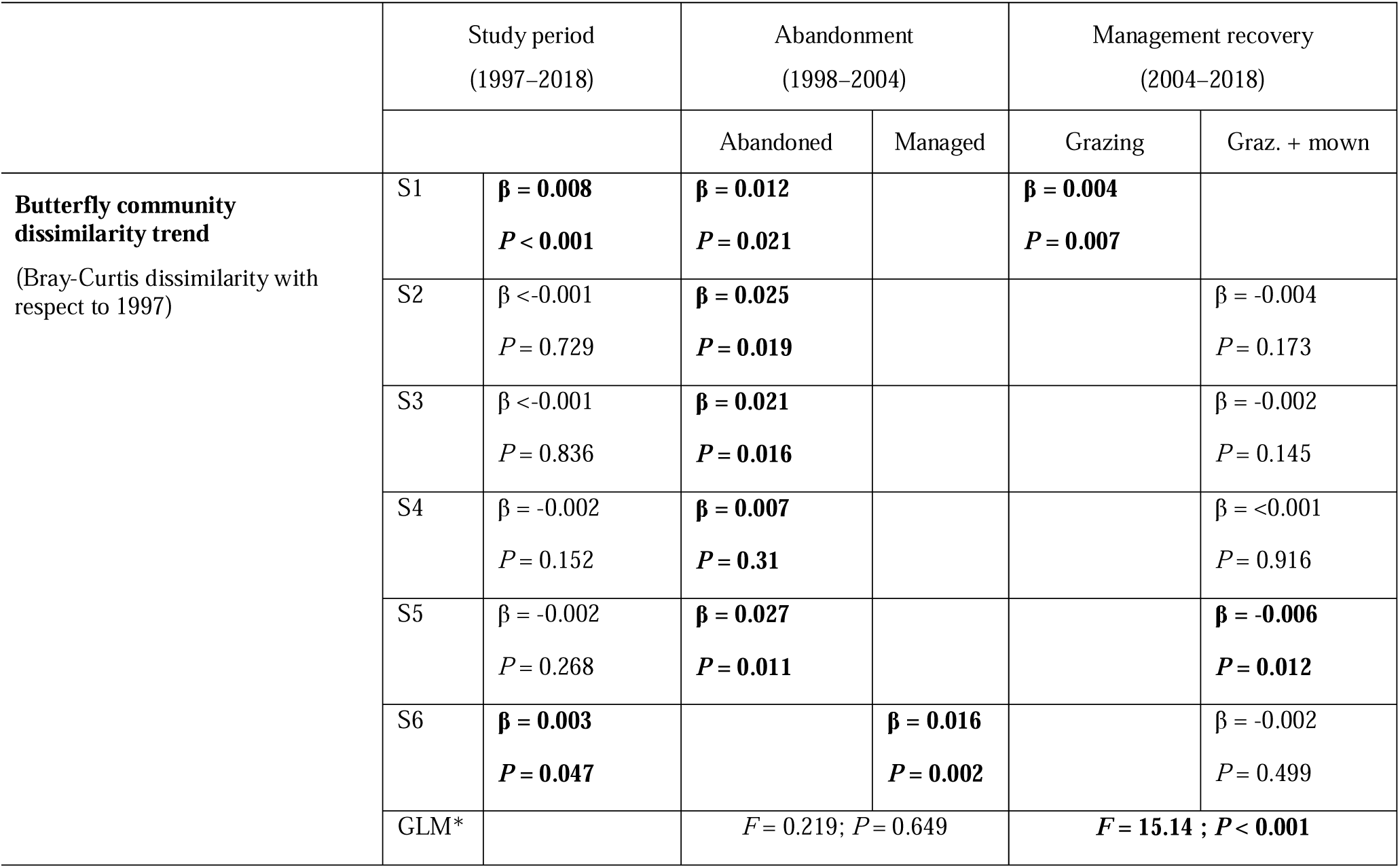
Butterfly community dissimilarity trends in the analysed periods. Beta coefficients and *P* valus are given

**Table A2.** PERMANOVA results for the butterfly community between periods: (A): before abandonment*; (B) abandonment; (C) management recovery. * 1999 was included in this period as butterfly communities were still very similar to the initial situation due to the inertia in changes in plant composition in the first year after abandonment (see Figure 6).

| Section | Treatment | PERMANOVA analysis | A vs B | A vs C | B vs C |
| --- | --- | --- | --- | --- | --- |
| 1 | Abandoned<br>Only-grazed | *P* < **0.001** | *P* = **0.045** | *P* = **0.005** | *P* < **0.001** |
| 2 | Abandoned<br>Mowing-grazing | *P* = **0.009** | *P* = 0.158 | *P* = 1 | *P* = **0.015** |
| 3 | Abandoned<br>Mowing-grazing | *P* = **0.001** | *P* = 0.096 | *P* = 0.334 | *P* = **0.001** |
| 4 | Abandoned<br>Mowing-grazing | *P* < **0.001** | *P* = 0.164 | *P* = 0.052 | *P* = **0.001** |
| 5 | Abandoned<br>Mowing-grazing | *P* = **0.001** | *P* = 0.052 | *P* = 1 | *P* < **0.001** |
| 6 | Managed<br>Mowing-grazing | *P* = **0.018** | *P* = 0.105 | *P* = 0.084 | *P* = **0.006** |

**Figure A1.**
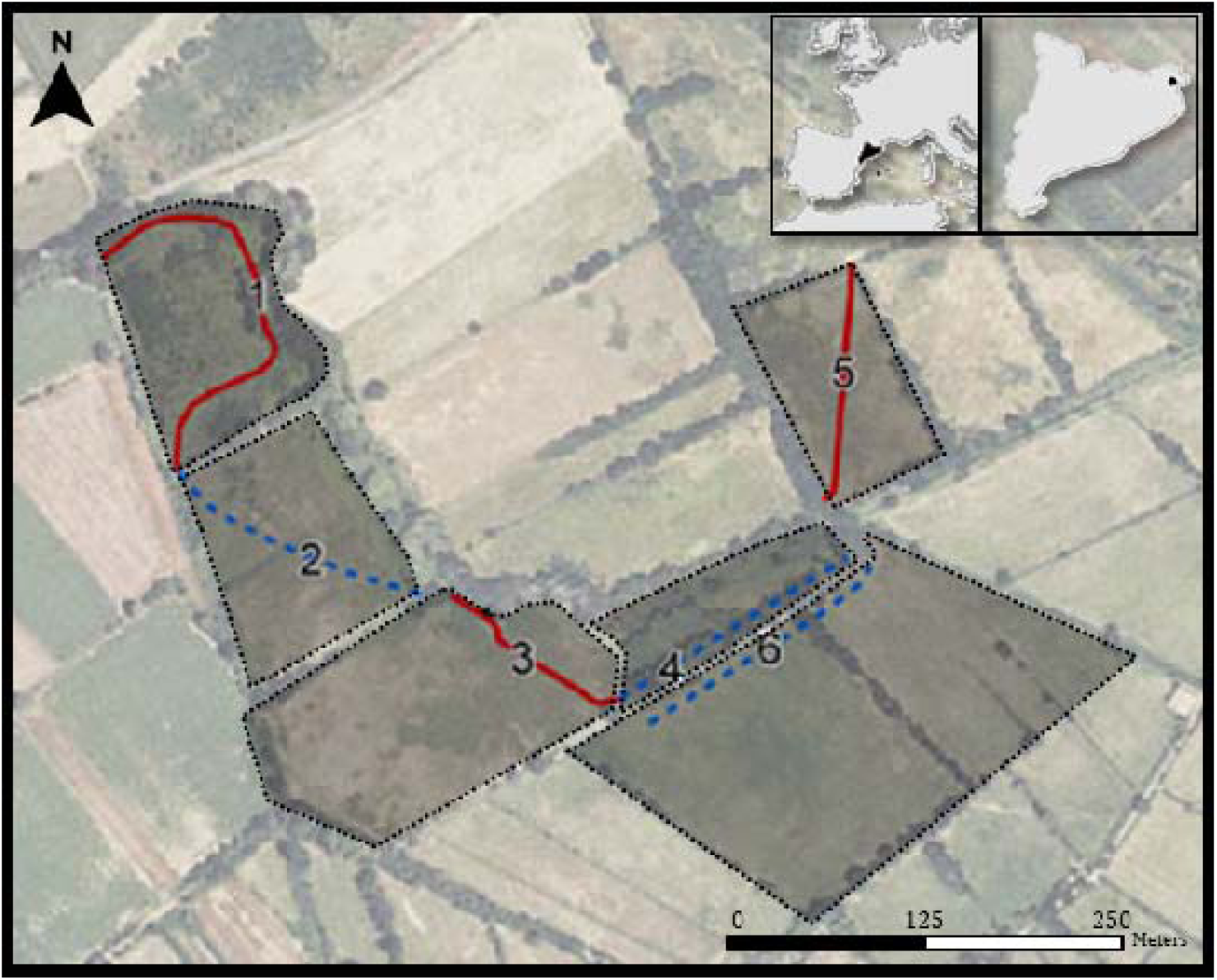
Study area. Transect sections in Closes de Tec, Aiguamolls de l’Empordà Natural Park.

**Figure A2.**
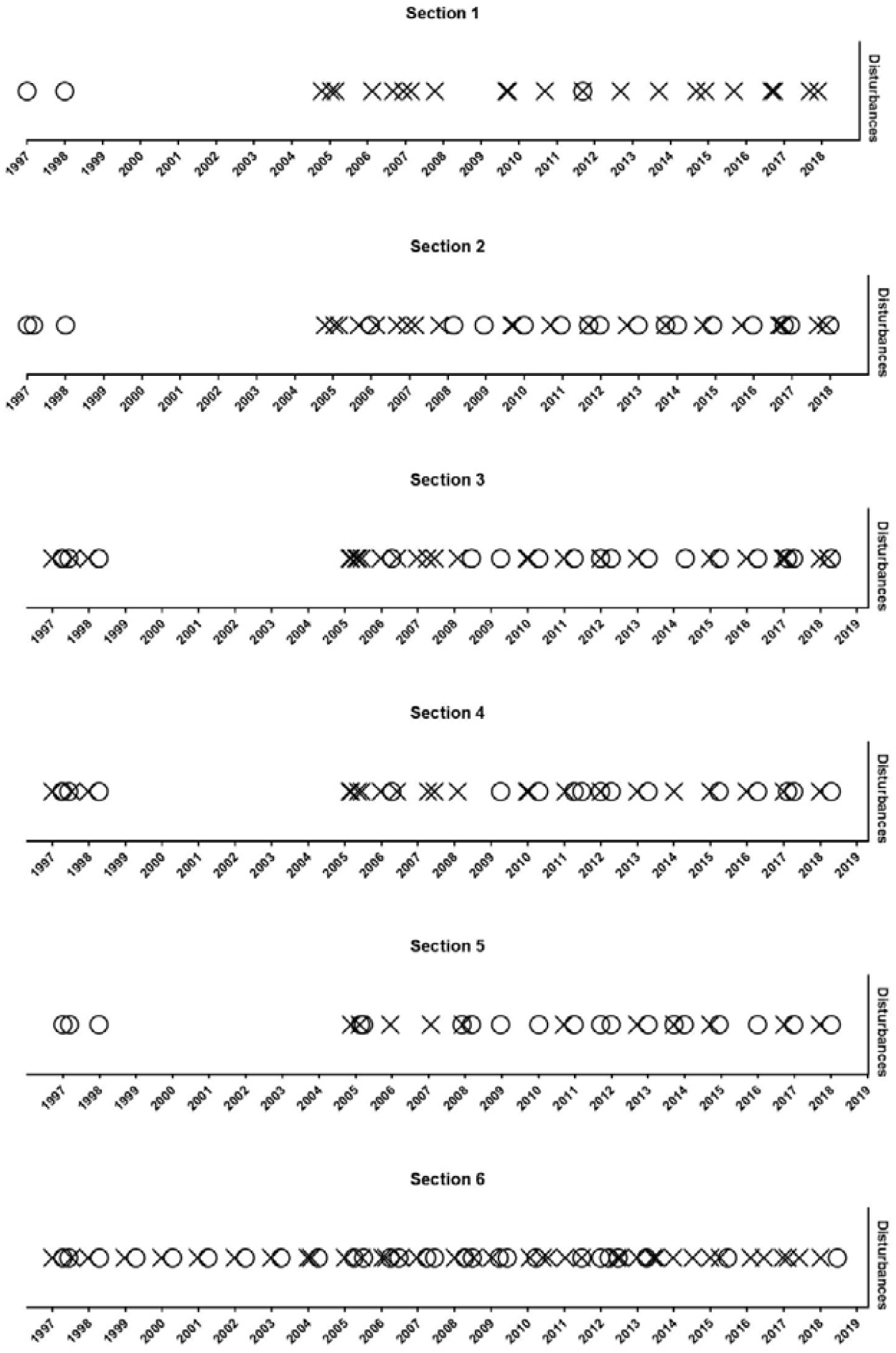
Type of management in the sections during the study period. Circles indicate episodes of grazing and crosses indicate mowing. As of 1999, sections 1–5 were abandoned; section 6 continued to be managed throughout the whole study period. Management was restored in 2005 in the abandoned meadows, although, while sections 2–5 combined grazing and mowing, section 1 was managed exclusively by grazing.

**Figure A3.**
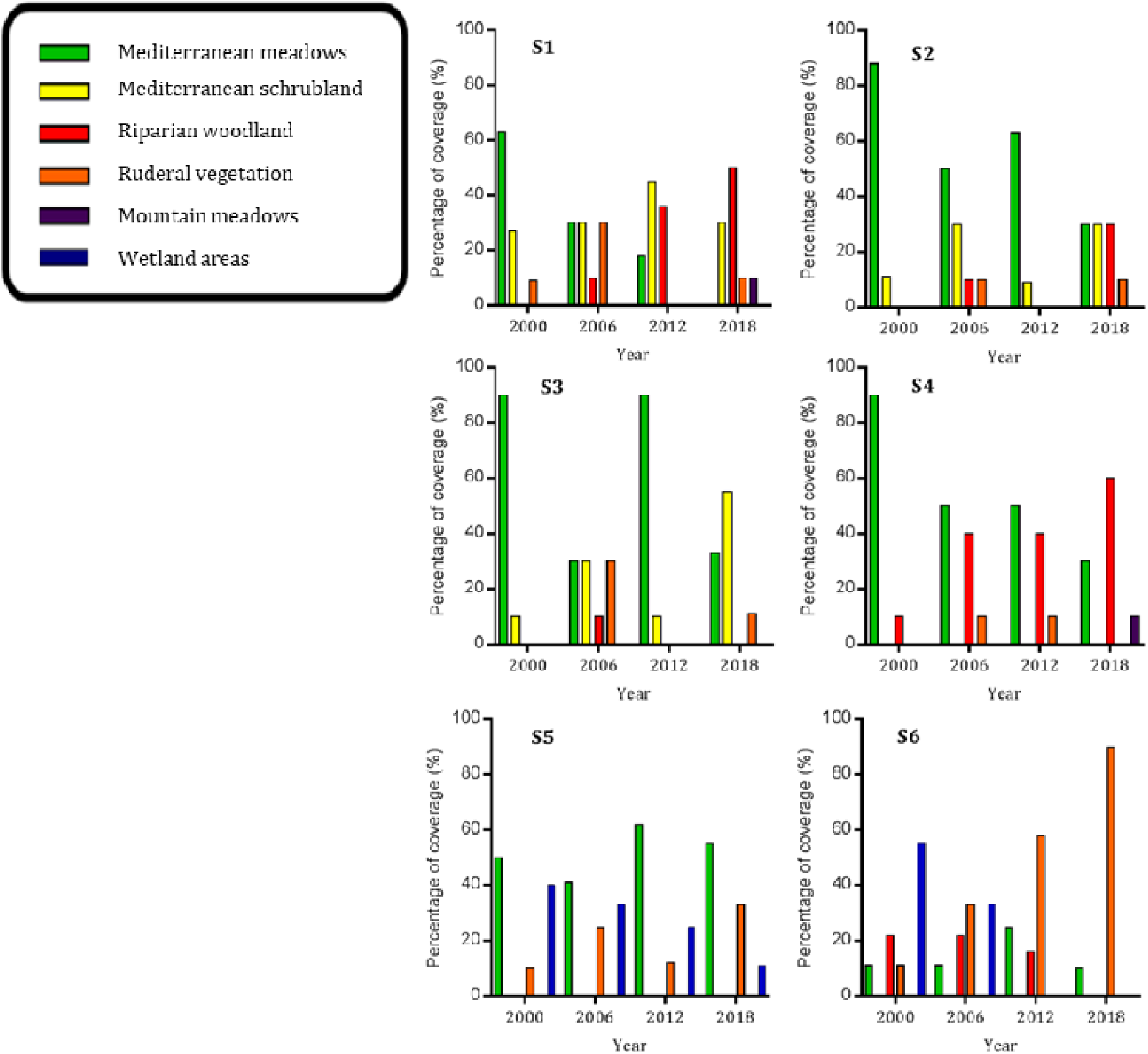
Habitat cover present along the different sections in 2000, 2006, 2012 and 2018. Habitat types characterized according to the CORINE Land Cover manual.

**Figure A4.**
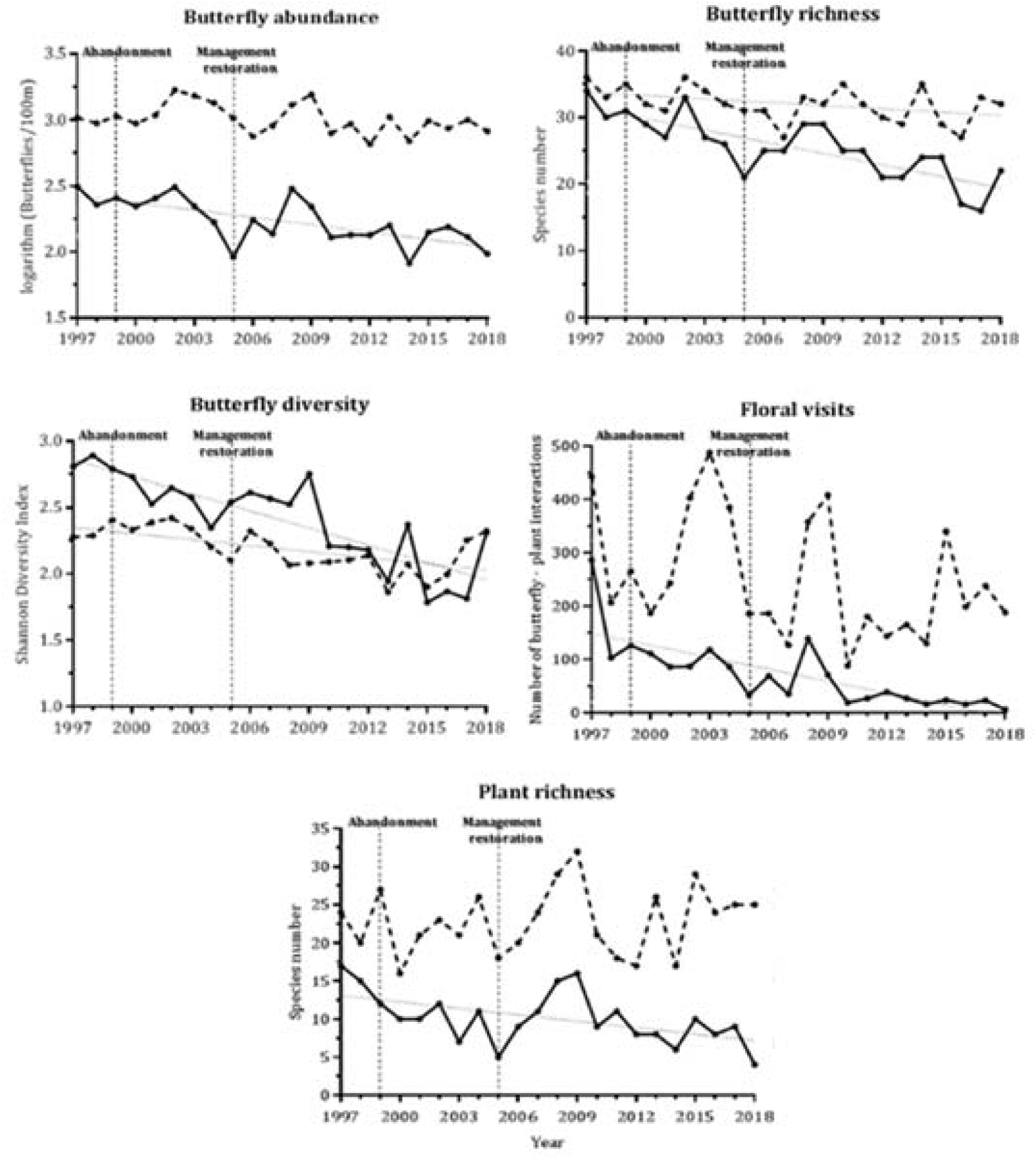
Continuous lines indicate trends for the eight ecological descriptors analysed in section 1. Dashed lines indicate trends for the total of all six sections. Significant trends (*P* < 0.05) are represented by small dotted lines.

**Figure A5.**
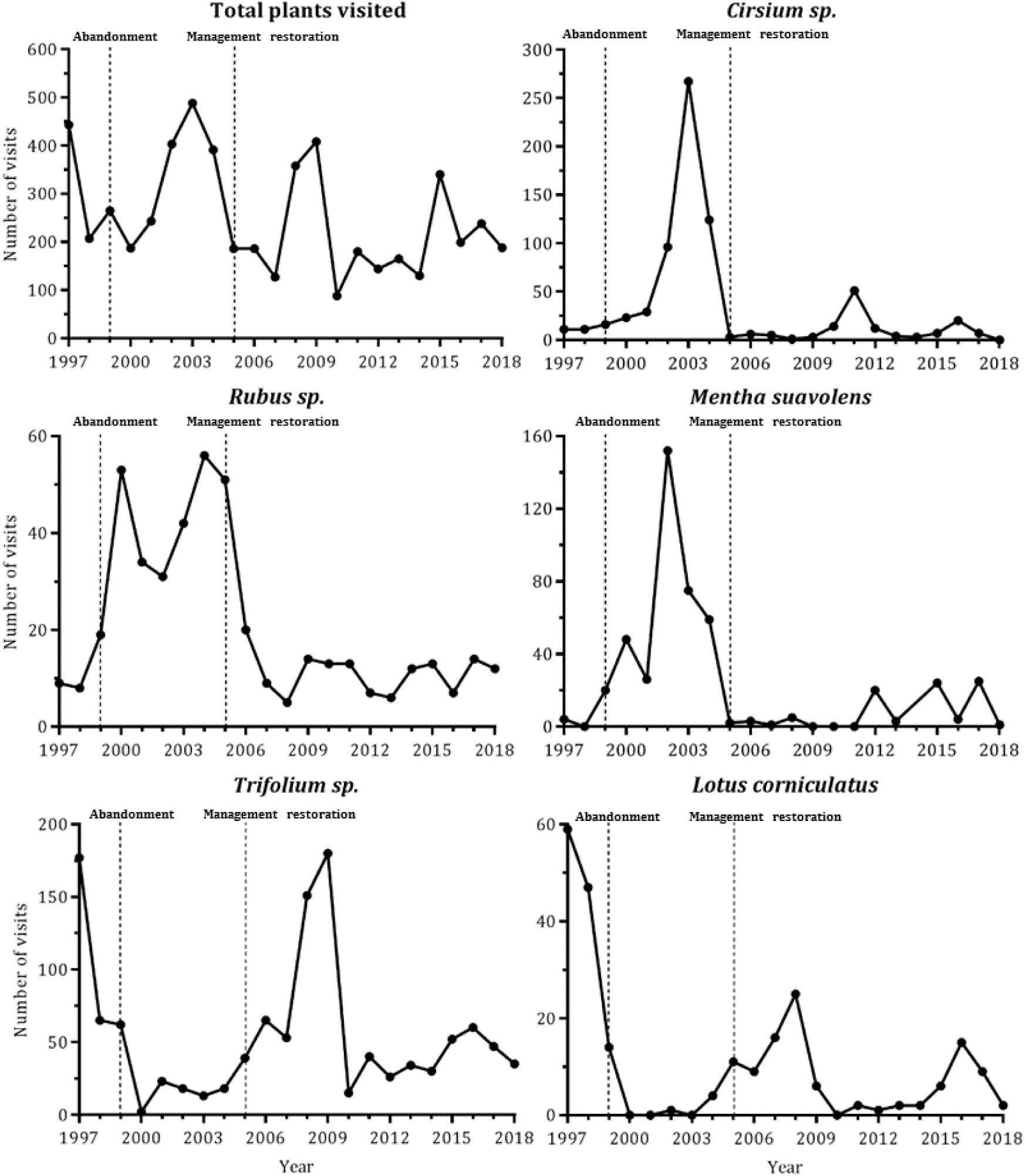
Population trends of the most sensitive plant species to management changes according to a Simper analysis. Trends for the totals for the five abandoned sections.

**Figure A6.**
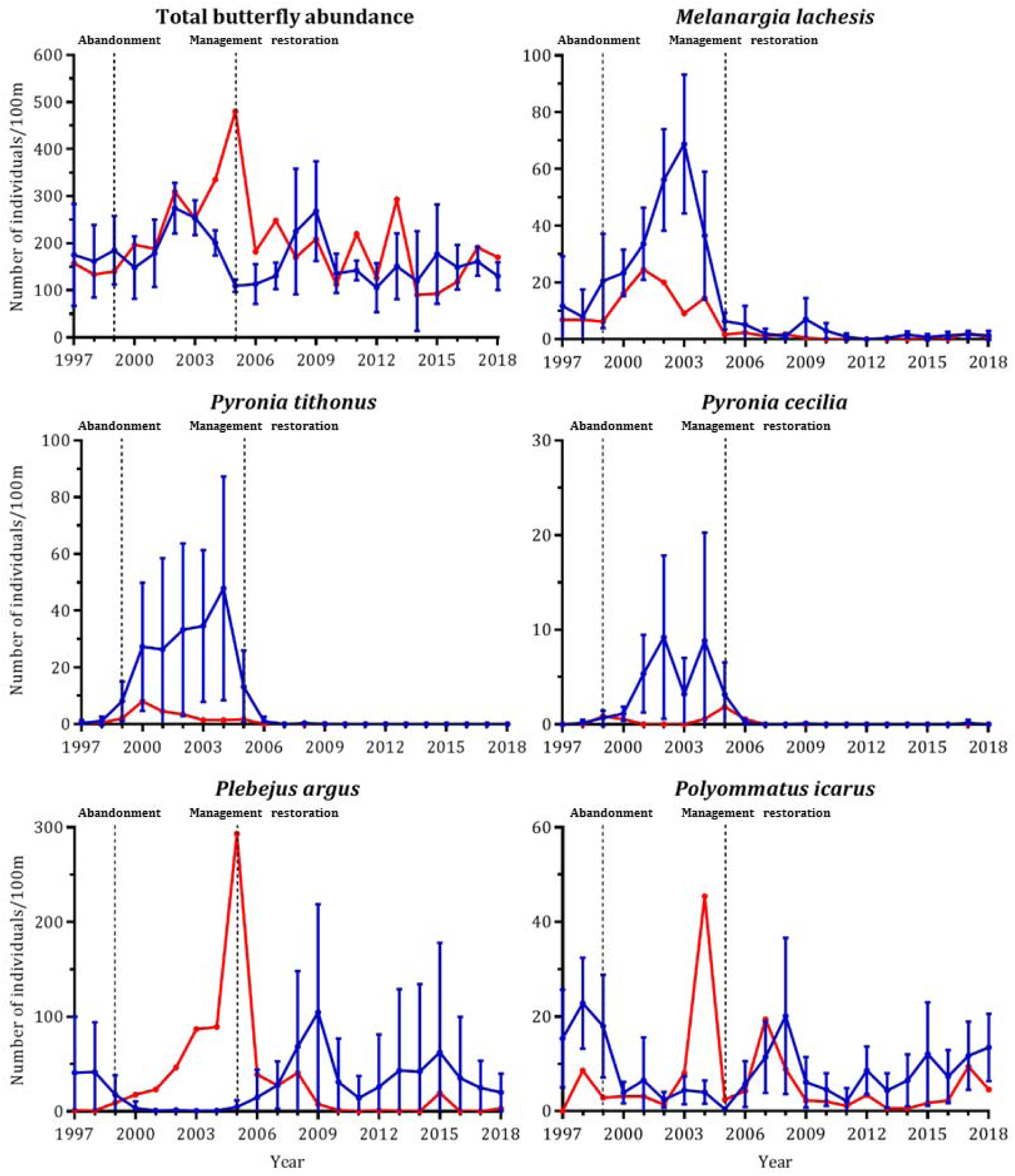
Population trends of the most sensitive butterfly species to management changes according to a Simper analysis. Blue lines show the trends in species in abandoned sections (1–5) and red lines trends in species in section 6.

